# Neonatal Resting-State Functional Connectivity Predicts Socioemotional and Behavioral Outcomes at 18 Months

**DOI:** 10.64898/2026.04.21.719787

**Authors:** Mi Zou, Arun L. W. Bokde

## Abstract

Early behavioral and temperamental differences are important indicators of later socioemotional development and psychopathology risk, yet their neural bases near birth remain incompletely understood. Using resting-state fMRI data from the Developing Human Connectome Project, we examined whether neonatal functional connectivity predicts 18-month behavioral and temperament outcomes in 397 infants (277 term-born, 120 preterm-born). Outcomes were assessed using the Child Behavior Checklist (CBCL) and the Early Childhood Behavior Questionnaire (ECBQ). We applied a stability-driven, ROI-constrained connectome-based predictive modeling framework to identify robust whole-brain connectivity features associated with measures of externalizing, internalizing, surgency, negative affect, and effortful control at 18 months. Statistically significant predictive models were observed for multiple outcomes across the whole cohort as well as within term-born and preterm-born groups, with clear differences in brain networks between cohorts. Across analyses, prefrontal and temporoparietal regions were repeatedly implicated, alongside medial temporal, fusiform, parahippocampal, and orbitofrontal-related regions. These findings indicate that large-scale neonatal functional organization is meaningfully associated to later socioemotional and behavioral differences, and that preterm birth is associated with differences in which brain networks are associated with later behaviour.

## Introduction

Early behavioral and temperamental differences are important markers of later socioemotional development and psychopathology risk. Dimensions such as externalizing behavior, internalizing behavior, surgency, negative affect, and effortful control capture early individual differences in emotional reactivity, regulation, and adaptation that may indicate later mental health outcomes (Kelsey et al., 2021; Bäuml et al., 2019). Identifying neural correlates of these phenotypes near birth may therefore help clarify how risk develops in the developing brain.

Resting-state functional MRI (rs-fMRI) has shown that the neonatal brain already exhibits organized large-scale functional architecture, providing a basis for studying how early connectivity relates to later development (Fransson et al., 2007). A growing number of studies indicate that neonatal functional connectivity is associated with affective and behavioral differences. For example, neonatal connectivity within emotion-related circuits has been linked to early-emerging callous-unemotional traits, fear-related behavior, negative reactive temperament, and internalizing symptoms (Brady et al., 2024; Thomas et al., 2019; Filippi et al., 2021; Rogers et al., 2017). More broadly, variability in infant functional connectivity has been associated with differences in affect and behavior, and infant frontoparietal and default mode connectivity has been linked concurrently and prospectively to negative affectivity (Kelsey et al., 2021; Ravi et al., 2023). Together, these findings suggest that early functional organization is already meaningfully related to later socioemotional functioning.

Most prior neonatal rs-fMRI studies, however, have focused on predefined circuits or seed-based analyses, particularly amygdala-centered pathways, rather than asking which whole-brain networks carry the most stable and generalizable predictive signal. Although previous work has provided important insight into early affective circuitry, it leaves open whether broader functional systems beyond canonical limbic pathways contribute to later internalizing, externalizing, and temperament outcomes. In addition, prior studies have often examined one outcome domain at a time rather than testing multiple related behavioral dimensions within a common modeling framework (Filippi et al., 2021; Ramphal et al., 2020; Ravi et al., 2023). Much of the literature also remains association-based, limiting inference about which connectivity patterns may support individual-level prediction.

These gaps may be especially important in pre-term neonates. Preterm birth occurs during a period of rapid network maturation, and extrauterine exposure during late gestation may alter neural development in ways relevant to later affective and behavioral outcomes. Very early preterm children show atypical resting-state neuromagnetic connectivity, and third-trimester extrauterine exposure has been linked to altered development of central autonomic network connectivity (Kozhemiako et al., 2019; Christoffel et al., 2025). Because early behavioral regulation and emotional reactivity depend on coordination across sensory, affective systems, the neural architecture predicting later affective and behavioral outcomes may differ between term-born and preterm-born infants.

Using data from 402 infants in the Developing Human Connectome Project (278 term-born and 124 preterm-born), we applied a region-of-interest (ROI)-constrained variant of Connectome-Based Predictive Modeling (CPM) to predict 18-month socioemotional and behavioral outcomes measured using the Child Behavior Checklist (CBCL) and the Early Childhood Behavior Questionnaire (ECBQ). Our approach extends standard CPM by introducing a hub-oriented framework that identifies high-degree ROIs from stable edge selection and restricts predictive feature selection to edges connected to these ROIs. This design was motivated by the possibility that predictive performance in neonatal rs-fMRI is weakened when low-signal or unstable features are included indiscriminately. We therefore aimed to identify robust and interpretable whole-brain connectivity signatures of later behavioral and temperament outcomes, and to test whether their predictive brain networks differs between term-born and preterm-born infants.

## Methods

### Participants

We analyzed data from 397 participants selected from the 814 available functional MRI (fMRI) datasets in the Developing Human Connectome Project(dHCP, Release 4). Among 124 preterm-born infants, 91 had two imaging sessions; in such cases, only the first session, scanned shortly after birth, was included in the analysis. Participants were excluded based on the following criteria: (i) excessive head motion (n = 151; motion exclusion criteria see below), (ii) missing CBCL or ECBQ scores at 18 months (n = 121), and (iii) term-born infants with postnatal age (scan age - gestational age) >3 weeks (n = 54). The final cohort comprised 277 term-born and 120 preterm-born infants, with preterm defined as gestational age <37 weeks. Demographic information is presented in Table 1. Neurodevelopmental outcomes were assessed at 18 months’ corrected age using the Child Behavior Checklist (CBCL) and the Early Childhood Behavior Questionnaire (ECBQ).

**Table 1.** Research participants.

|  | Term(n=277) | Preterm(n=120) | Statistics | *p* value |
| --- | --- | --- | --- | --- |
| Demographics |  |  |  |  |
| GA at birth,<br>median(range),weeks | 40.14 (37.0-42.7) | 33.29 (23.0-36.9) | 15.8<sup>a</sup> | p < 0.001 |
| PMA at scan,<br>median(range),weeks | 41.14 (37.4-44.7) | 35.41 (27.3-45.1) | 13.5<sup>a</sup> | p < 0.001 |
| Female(%) | 125 (45) | 51 (42) | 0.2<sup>b</sup> | 0.629 |
| Weight at birth,<br>median(range),kg | 3.77 (2.7-4.3) | 1.64 (0.5-4.1) | 10.8<sup>a</sup> | p < 0.001 |
| % FD outliers(SD) | 4.25 (2.7) | 4.75 (2.4) | -0.5<sup>a</sup> | p = 0.650 |
| Follow-up |  |  |  |  |
| CBCL composite score,<br>mean(SD)<sup>c</sup> | 47.61 (8.9) | 46.57 (9.9) | 1.2<sup>a</sup> | p = 0.243 |
| Internalizing score<sup>c</sup> | 44.72 (9.7) | 44.71 (10.5) | 0.0<sup>a</sup> | p = 0.973 |
| Externalizing score<sup>c</sup> | 48.28 (8.7) | 46.98 (9.5) | 1.5<sup>a</sup> | p = 0.141 |
| Surgency score, mean(SD)<sup>d</sup> | 5.21 (0.7) | 5.28 (0.9) | -1.2<sup>a</sup> | p = 0.220 |
| Negative affect score<sup>d</sup> | 2.68 (0.7) | 2.60 (0.8) | 1.5<sup>a</sup> | p = 0.138 |
| Effort control score<sup>d</sup> | 4.51 (0.7) | 4.66 (0.7) | -1.9<sup>a</sup> | p = 0.060 |

FD framewise displacement <sup>a</sup>Z(Mann-Whiteney U test) <sup>b</sup> $\chi^2$ -test <sup>c</sup>Child Behavior Checklist <sup>d</sup>Early Childhood Behavior Questionnaire

### MRI Data Acquisition

Functional MRI data were collected as part of the developing Human Connectome Project (dHCP) at the Evelina Newborn Imaging Centre, Evelina London Children’s Hospital, utilizing a 3 Tesla Philips Achieva system. The study received ethical clearance from the UK National Research Ethics Authority (14/LO/1169), and all participating families provided written informed consent prior to the imaging sessions. All scans were performed without the use of sedation in a neonatal environment designed specifically for safety and comfort, which included a custom 32-channel head coil, an acoustic hood, and devices to ensure proper positioning of the infants. Infants wore MRI-compatible ear protection to reduce noise exposure, and a neonatal nurse or paediatrician continuously monitored vital signs, including heart rate, oxygen saturation, and body temperature.

Blood-oxygen-level-dependent (BOLD) fMRI data were acquired using a multi-slice echo planar imaging sequence with multiband excitation (factor 9) over a scan duration of 15 minutes and 3 seconds, producing 2300 volumes. Imaging parameters included a repetition time (TR) of 392 ms, echo time (TE) of 38 ms, a flip angle of 34°, and a voxel resolution of 2.15 mm isotropic. High-resolution T2w anatomical images were obtained for structural analysis and functional data registration. T2-weighted images were obtained with a resolution of 0.8 mm isotropic and a field of view of 145 × 145 × 108 mm, a TR of 12 s, and a TE of 156 ms.

### Functional data preprocessing

The preprocessing of neuroimaging data was performed using a bespoke pipeline specifically optimized for neonatal imaging and developed for the dHCP, as detailed in Fitzgibbon et al. (2020). The pipeline accounted for susceptibility-induced dynamic distortions as well as intra- and inter-volume motion artifacts. Twenty-four extended rigid-body motion parameters were regressed alongside single-subject independent component analysis (ICA) noise components identified using the FSL FIX tool (Oxford Centre for Functional Magnetic Resonance Imaging of the Brain’s Software Library, version 5.0). The denoised data were first registered to the T2-weighted native space using boundary-based registration. They were then non-linearly transformed to a standard space with a weekly template from the dHCP volumetric atlas through diffeomorphic multimodal (T1/T2) registration.

Because head motion is a potential surrogate marker for the infant’s arousal state and can interact with underlying neural activity, we adopted a conservative approach to minimize the impact of motion-related artifacts. Specifically, for each subject, we selected a continuous subset (1600 from the original 2300 acquired volumes) with the minimum total framewise displacement (FD) and the dataset was cropped accordingly – the cropped subset was used for all subsequent analyses. After the cropping, volumes were flagged as motion outliers based on FD, with outliers identified as FD >1.5 interquartile range (IQR) above the 75th centile. Subjects with more than 160 motion-outlier volumes (>10% of data) were excluded from further analyses. The motion outliers of the resulting data from the participants are showed in Table 1. No significant differences were observed between term and preterm groups under the assumption of non-normality (p = 0.650, Mann-Whitney U-test). The number of outliers was included as a covariate in subsequent regression analyses.

### Connectivity Matrices

Whole-brain functional connectivity was computed using the CONN toolbox (Whitfield-Gabrieli & Nieto-Castanon, 2012). Brain regions were defined using the Schaefer 200-node cortical atlas (Schaefer et al., 2018) combined with 8 subcortical regions from the dHCP parcellation, yielding 208 nodes. The atlas was aligned to the 40-week template via rigid-body, affine, and SyN diffeomorphic transformations implemented in ANTs. Mean time courses were extracted for each ROI, and pairwise Pearson correlations were computed and Fisher Z-transformed, resulting in a symmetric 208×208 connectivity matrix per subject.

### ROI-constrained Connectome–Based Predictive Modeling (R-CPM)

The pipeline was applied separately to preterm-only, term-only, and whole cohort (term + preterm), with preterm status included as a covariate in the whole cohort analysis. All analyses are performed separately for positive and negative associations. The full procedure is described here for convenience of reader; method was initially described in Zou & Bokde (Zou & Bokde, 2026).

#### Stage 1 — Stable edge discovery

We ran 10-fold repeated 150 times with new partitions. For each fold we computed partial correlations between each edge and behaviour controlling for gender, head motion (number of outliers), age at scan, and preterm status when applicable. Edges with r>0 and p<0.05 were counted as positive; r<0 and p<0.05 as negative. Counts were accumulated across all folds and repeats. An edge was labelled stable if selected in ≥95% of the 10×150 iterations. ROI ranking(below) was performed separately for positive and negative graphs.

#### Stage 2 — ROI ranking

From the Stage-1 stable graph (sign-specific), we identified hubs by iteratively removing the highest-degree ROI and recomputing degrees on the remaining graph. The ROI with the largest degree was labelled the top hub; after removing it and its incident edges, the ROI with the largest updated degree was labelled second, and so on, yielding a ROI order *R* = (*r*_1_, *r*_2_, …) (Figure 1a). No Stage-1 edges were carried forward as features; only the ROI order *R* was passed to Stage 3.

**Figure 1.**
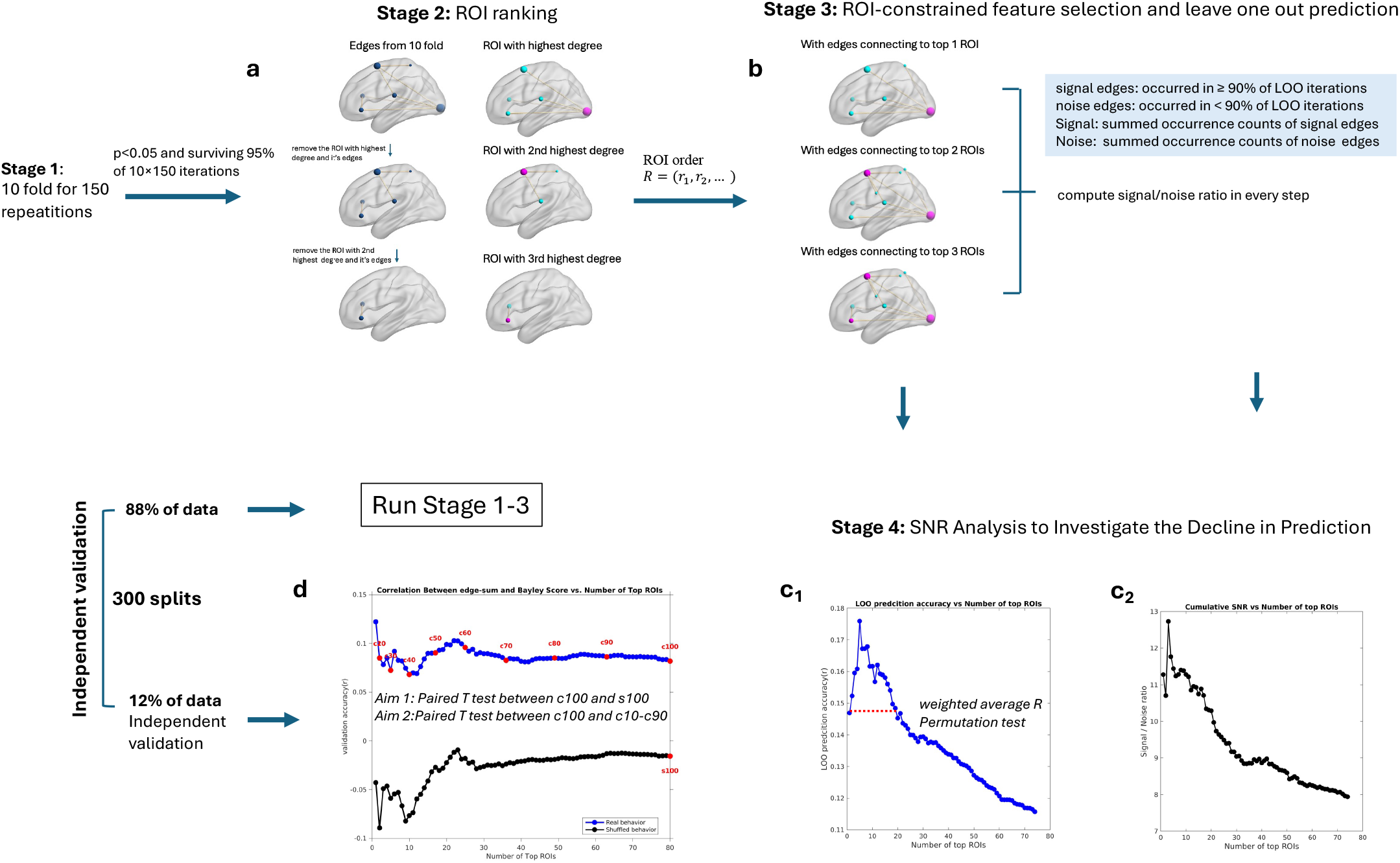
ROI-constrained CPM pipeline with ROI-constrained feature selection and independent validation. (a) Stage-2 — ROI ranking: ROIs were sorted by iteratively removing the highest-degree ROI to define a high-to-low ROI degree order. No Stage-1 edges were carried forward as features; only the ROI order *R* was passed to Stage 3. (b) Stage-3 — ROI-constrained CPM : For each ROI-set size (n = 1…|R|), edges that survived p<0.05 and connected to at least one of the top-n ROIs were selected via LOO to calculate prediction accuracy. SNR was also computed for each prefix n by dividing the summed occurrence counts of signal edges (occurred in ≥ 90% of LOO iterations) by those of noise edges (occurred in ≥ 90% of LOO iterations);(c) Stage-4 — SNR analysis: As progressively lower-degree ROIs were added, SNR steadily decreased(c_2_), mirroring the decline in predictive accuracy(c_1_). And SNR and prediction accuracy were strongly correlated (r = 0.96, p < 0.0001). We conducted a permutation test to confirm that the observed weighted average prediction performance (weighted by degree of each ROI) across all ROI-set sizes n exceeded chance; (d) Independent validation: For each ROI-set size (n = 1…|R|), validation subjects’ network strength—sums of signal edges incident on the top-n ROIs—were correlated with behavior to yield r curves. The cumulative distribution of edge counts across ROIs was computed, and thresholds were set at 10 %, 20 %, …, 100 % coverage of total edges. Aim 1 — Check whether all signal edges contribute predictive information: Paired T test between observed and shuffled behavior at coverage 100%. Aim 2 — Check whether the predictive effect declines as signal edges of low degree ROIs are added: Paired t-tests compared performance at each intermediate coverage level (10–90 %) with that at 100 %. Abbreviations: c10, coverage 10%; s100, shuffled behavior at coverage 100%; ROI, region of interest; LOO, leave one out; SNR, signal to noise ratio.

#### Stage 3 — ROI-constrained feature selection and leave-one-out (LOO) prediction

We adopted a ROI-constrained variant of CPM. For each prefix size *n* = 1, …, | *R* |(looping over the ordered ROIs) and each sign, we performed LOO: (1) Feature selection: Within each LOO training set (excluding the held-out subject), we computed partial correlation between each edge and behaviour(the same covariates applied), retaining edges meeting p<0.05 that connected to at least one of the top-*n* ROIs (Figure 1b). (2) Summary edges: For each subject computed *S*^+^ (sum of positive selected edges) and *S*^-^(sum of negative selected edges). (3) Prediction: We fitted separate linear models on the LOO training set for *S*^+^and *S*^-^and predicted behaviour of the left-out subject. (4) Edge stability across LOO: After LOO completed for a given *n*, retaining edges occurring in ≥90% of LOO iterations as 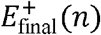 and 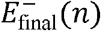. Stage 3 yielded, for each *n*, sign-specific LOO accuracy (Figure 1c1) and training-stable edge sets. As figure 1c1 shows, as the number of top ROIs increases, the prediction accuracy goes down.

#### Stage 4 —Signal to noise ratio (SNR) analysis to investigate the decline in prediction

To investigate why the predictive accuracy decreased as more ROIs were added, we examined how the inclusion of lower-degree ROIs affected the SNR.

For each prefix size *n* = 1,…,|R| (looping over the ordered ROIs), we recorded occurrence of all edges across the LOO iterations in Stage 3. These edges were divided into two categories based on a stability threshold of 0.9 × (number of LOO iterations): (1) Signal edges: edges with occurrence counts ≥ threshold; (2) Noise edges: edges with occurrence counts < threshold.

The SNR for each prefix *n* was then computed as

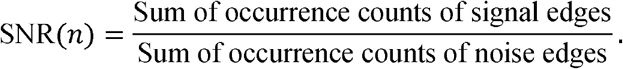

As progressively lower-degree ROIs were added in the whole-cohort model of composite cognition, the SNR consistently declined (Figure 1c2). And SNR was strongly correlated with LOO prediction accuracy (r = 0.96, p < 0.0001; Figures 1c1–1c2). Similar relationships between SNR and prediction accuracy were observed across all behaviours and cohorts (Supplementary Table 1), indicating that the decrease in prediction arises from a reduced SNR as low degree ROIs are incorporated.

### Permutation testing of prediction accuracy

After demonstrating that the decline in predictive accuracy with increasing number of top ROIs reflects a reduction in SNR, we conducted a permutation test to confirm that the observed weighted average prediction performance (weighted by degree of each ROI) across all ROI-set sizes *n* (figure 1c1) exceeded chance. Across Stages 1–3, behavioural labels were randomly permuted while preserving the full analytical pipeline. Because CBCL and ECBQ scores in this healthy neonatal cohort included many with same values, we excluded permutations in which >40% of the scores were reassigned to their original positions. In cases where no edges survived across the 10 folds in Stage 1, the corresponding prediction performance was assigned a value of zero. For each permutation, we computed the weighted average R across all ROI-set sizes *n*. The one-sided *p*-value was calculated as following:

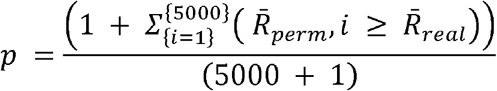

5000= total number of permutations. *R*_perm,*i*_ = weighted average correlation between predicted and permuted behavioral scores across all ROI set sizes *n. R*_real_ = weighted average correlation between predicted and observed behavioral scores across all ROI set sizes *n*.

### Independent validation of signal edge effects and ROI-dependent decline

We have established that the reduction in predictive accuracy is driven by the growing contribution of noise from edges originating in low-degree ROIs. It remains unclear whether all signal edges—including those connected to low-degree ROIs—contribute predictive information, and whether prediction accuracy decreases as signal edges from low-degree ROIs are added.

In order to investigate this, we conducted an independent validation analysis. Approximately 88% of the data were used for running stage 1-3, while the remaining 12% were reserved for independent validation. The split was repeated for 300 times. All feature discovery and model fitting used training data only; validation data were untouched until final testing.

To preserve the discretized outcome distribution, we stratified the split by exact behavioural score value: for values with ≥8 subjects we assigned ∼88% to training and ∼12% to validation; values with <8 subjects were pooled before applying the same proportion.

For each sign and each prefix size *n = 1,…*,|*R*| (looping over the ordered ROIs), we used the training-stable edge sets 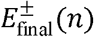 (signal edges from Stage 3): (1)For each validation subject, we computed *S*^+^/ *S*^-^by summing all edges incident on the top-*n* ROIs; (2)Each subject’s *S*^±^ was correlated (Pearson’s *r*) with behaviour to yield predictive-accuracy curves *r*^±^(*n*); (3) A single-shuffle was performed by permuting the behaviour once per split to estimate chance-level *r*^±^(*n*) values (Figure 1d).

#### Aim 1 — Do all signal edges contribute predictive information?

To evaluate whether the entire set of signal edges retains predictive power, we focused on the largest ROI set (*n =* |*R*|) and did paired *t*-tests(fisher-z transformed r value) between real behaviour and shuffled behaviour across 300 splits to test whether the final model’s correlation remained significantly above the single shuffle (Figure 1d).

#### Aim 2 — Does the predictive effect decline as signal edges from low degree ROIs are added?

To examine how increasing the ROI set impacts performance, we quantified accuracy as a function of cumulative edge coverage. For each behavioral measure, the cumulative distribution of edge counts across ROIs was computed, and thresholds were set at 10 %, 20 %, …, 100 % of total edges. For each coverage level, the corresponding predictive correlations *r*(*n*) were obtained. Across 300 splits, paired *t*-tests (Fisher-z transformed r value) compared performance at each intermediate coverage level (10–90 %) with that at 100 % (Figure 1d).

### Statistical evaluation and reporting

The following two statistical test significance will be reported: (1) permutation statistical test of average R across all ROI-set sizes *n*; (2) independent validation test on the largest ROI set (*n =* |*R*|). Pooled mean validation accuracy (r□) at *n* = *R* with 95% confidence interval will be reported. Significance versus the behaviour-shuffle null at *n* = *R* is tested with a paired t-test across splits; we report *t*(*df*), exact *p*, and Cohen’s *d*_z_(see Table 2). Correlations were Fisher-z transformed for inference and back-transformed for presentation.

**Table 2.**
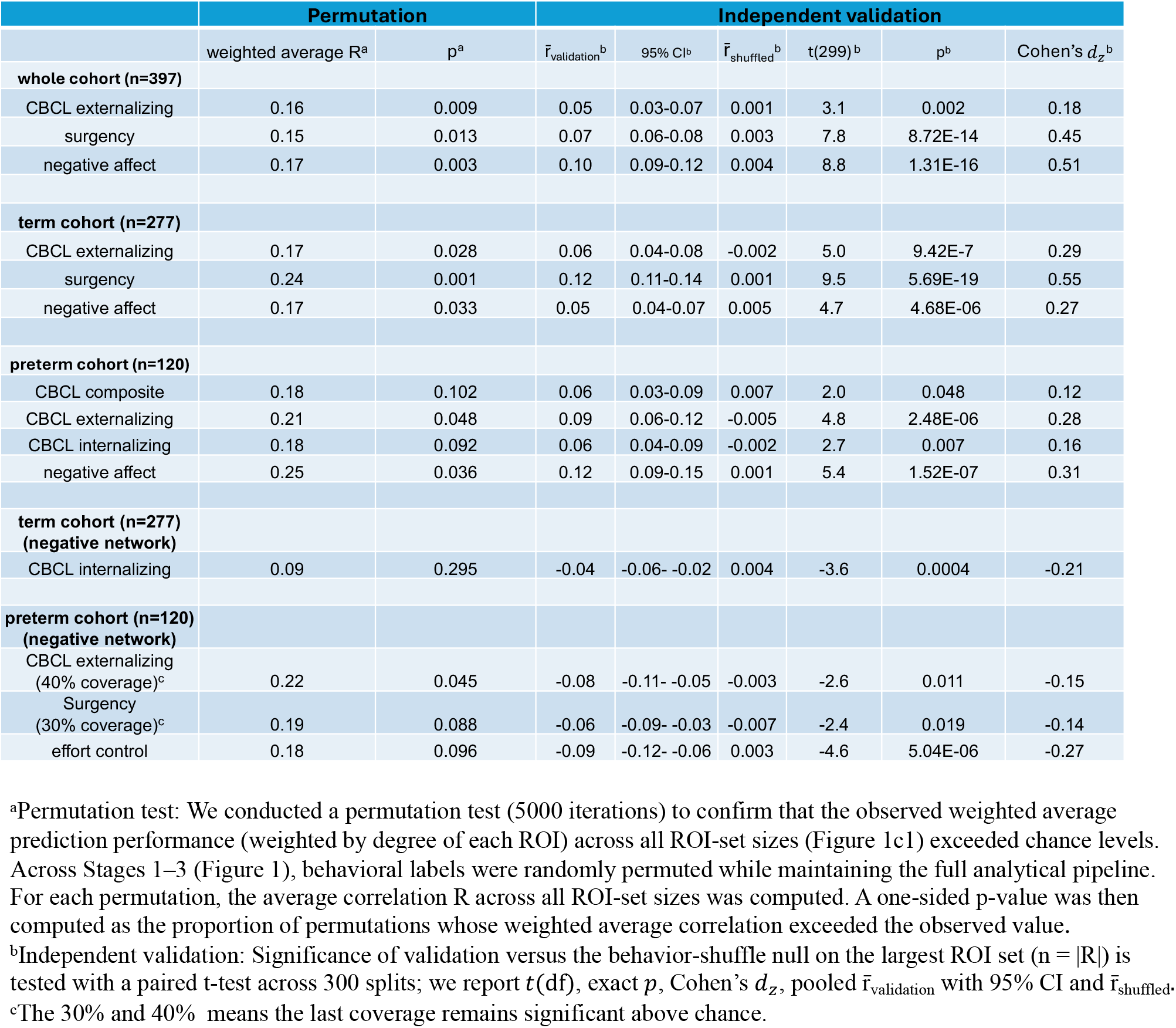
Statistics of permutation and independent validation.

|  | Permutation |  | Independent validation |  |  |  |  |  |
| --- | --- | --- | --- | --- | --- | --- | --- | --- |
| | weighted average R<sup>a</sup> | p<sup>a</sup> | $\bar{r}_{\text{validation}}^b$ | 95% CI<sup>b</sup> | $\bar{r}_{\text{shuffled}}^b$ | t(299)<sup>b</sup> | p<sup>b</sup> | Cohen's $d_z^b$ |
| **whole cohort (n=397)** |  |  |  |  |  |  |  |  |
| CBCL externalizing | 0.16 | 0.009 | 0.05 | 0.03-0.07 | 0.001 | 3.1 | 0.002 | 0.18 |
| surgency | 0.15 | 0.013 | 0.07 | 0.06-0.08 | 0.003 | 7.8 | 8.72E-14 | 0.45 |
| negative affect | 0.17 | 0.003 | 0.10 | 0.09-0.12 | 0.004 | 8.8 | 1.31E-16 | 0.51 |
| **term cohort (n=277)** |  |  |  |  |  |  |  |  |
| CBCL externalizing | 0.17 | 0.028 | 0.06 | 0.04-0.08 | -0.002 | 5.0 | 9.42E-7 | 0.29 |
| surgency | 0.24 | 0.001 | 0.12 | 0.11-0.14 | 0.001 | 9.5 | 5.69E-19 | 0.55 |
| negative affect | 0.17 | 0.033 | 0.05 | 0.04-0.07 | 0.005 | 4.7 | 4.68E-06 | 0.27 |
| **preterm cohort (n=120)** |  |  |  |  |  |  |  |  |
| CBCL composite | 0.18 | 0.102 | 0.06 | 0.03-0.09 | 0.007 | 2.0 | 0.048 | 0.12 |
| CBCL externalizing | 0.21 | 0.048 | 0.09 | 0.06-0.12 | -0.005 | 4.8 | 2.48E-06 | 0.28 |
| CBCL internalizing | 0.18 | 0.092 | 0.06 | 0.04-0.09 | -0.002 | 2.7 | 0.007 | 0.16 |
| negative affect | 0.25 | 0.036 | 0.12 | 0.09-0.15 | 0.001 | 5.4 | 1.52E-07 | 0.31 |
| **term cohort (n=277)**<br>**(negative network)** |  |  |  |  |  |  |  |  |
| CBCL internalizing | 0.09 | 0.295 | -0.04 | -0.06- -0.02 | 0.004 | -3.6 | 0.0004 | -0.21 |
| **preterm cohort (n=120)**<br>**(negative network)** |  |  |  |  |  |  |  |  |
| CBCL externalizing<br>(40% coverage)<sup>c</sup> | 0.22 | 0.045 | -0.08 | -0.11- -0.05 | -0.003 | -2.6 | 0.011 | -0.15 |
| Surgency<br>(30% coverage)<sup>c</sup> | 0.19 | 0.088 | -0.06 | -0.09- -0.03 | -0.007 | -2.4 | 0.019 | -0.14 |
| effort control | 0.18 | 0.096 | -0.09 | -0.12- -0.06 | 0.003 | -4.6 | 5.04E-06 | -0.27 |

<sup>a</sup>Permutation test: We conducted a permutation test (5000 iterations) to confirm that the observed weighted average prediction performance (weighted by degree of each ROI) across all ROI-set sizes (Figure 1c1) exceeded chance levels. Across Stages 1–3 (Figure 1), behavioral labels were randomly permuted while maintaining the full analytical pipeline. For each permutation, the average correlation R across all ROI-set sizes was computed. A one-sided p-value was then computed as the proportion of permutations whose weighted average correlation exceeded the observed value.
<sup>b</sup>Independent validation: Significance of validation versus the behavior-shuffle null on the largest ROI set ( $n = |R|$ ) is tested with a paired t-test across 300 splits; we report $t(df)$ , exact $p$ , Cohen's $d_z$ , pooled $\bar{r}_{\text{validation}}$ with 95% CI and $\bar{r}_{\text{shuffled}}$ .
<sup>c</sup>The 30% and 40% means the last coverage remains significant above chance.

We considered two cases. First, when performance at 100% coverage was significantly above chance, we further asked whether 100% coverage also yielded the best validation accuracy compared to other coverages. In most behaviors, the highest validation accuracy was observed at 100% coverage, indicating that predictive performance generally improved as signal edges associated with low-degree ROIs were added. In this case, we reported the signal edges that survived in 90% of splits at 100% coverage. Second, for behaviors in which the best coverage was below 100%, this was typically because a small subset of top ROIs accounted for a disproportionately large share of the total edges (see Supplementary Table 1 and Supplementary Figures S1). If 100 % coverage is still significant above chance, we still reported the signal edges that survived in 90% of splits at 100% coverage(we compared the connectivity patterns at the best coverage and at 100% coverage and found them to be consistent). If performance at 100% coverage was not significantly above chance, we report the signal edges that survived in 90% of splits at the last coverage level that remained significantly above chance. In both cases, all other tested coverage levels before the coverage we report also showed significant differences between real and shuffled data (paired t-test, p < 0.05).

The connectivity results are presented in a network format. Resting-state networks were defined using group-level independent component analysis of term-born infants scanned at 43.5–44.5 weeks postmenstrual age in the developing Human Connectome Project (dHCP) dataset (Eyre et al., 2021). Each ROI from the parcellation was assigned to the network with the highest spatial overlap (winner-takes-all criterion). The network map is shown in supplementary Figure S2.

## Results

The CBCL prediction model with the whole cohort was statistically significant for the CBCL externalizing score (see Table 2). The two dominant nodes are left fusiform gyrus and right medial temporal pole (Figure 3A), with temporoparietal network most actively involved, especially the connection between temporoparietal network and prefrontal network (Figure 4A).

**Figure 3.**
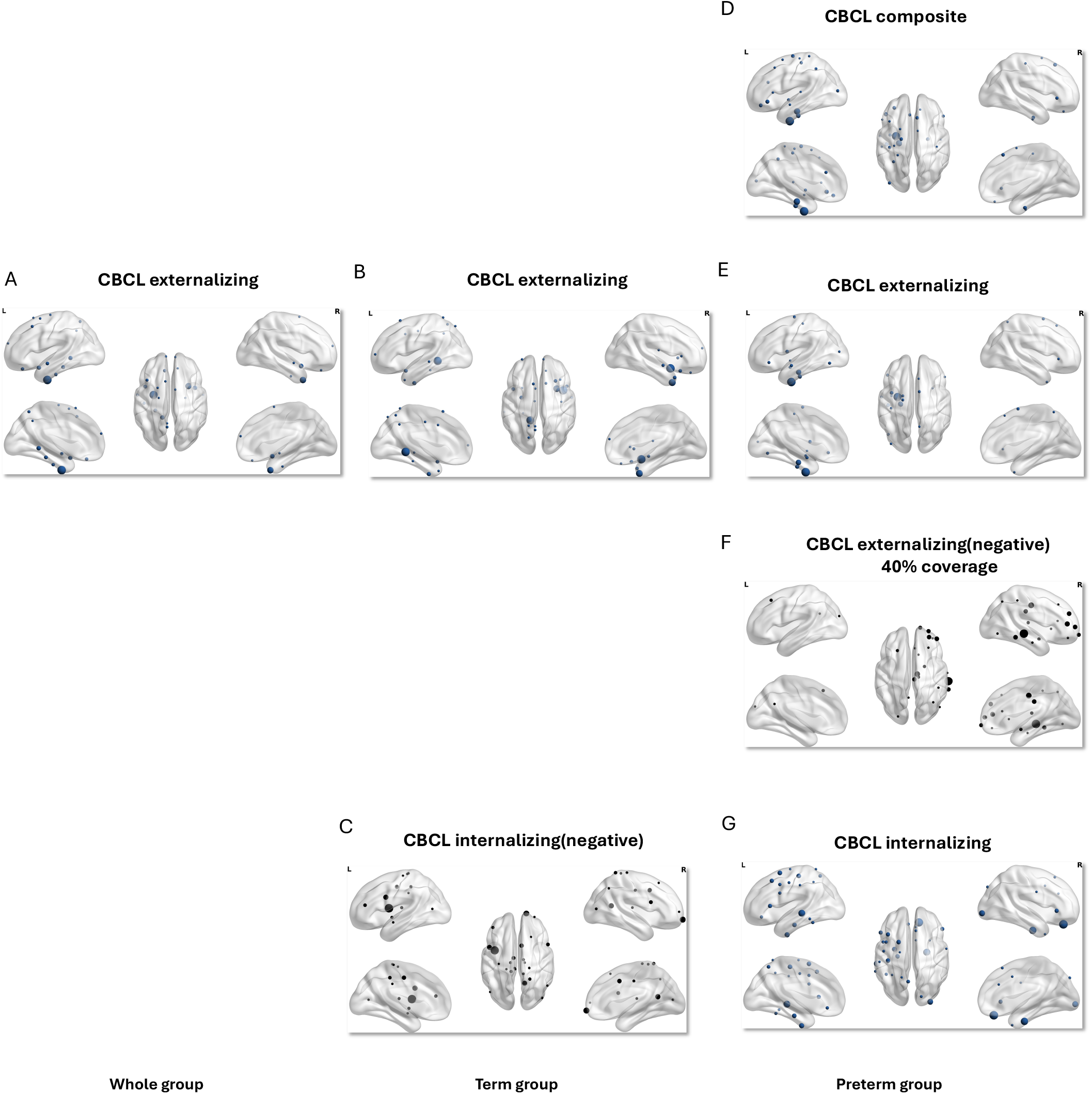
Predictive ROIs for CBCL: high-degree regions across cohorts and subscales. High-connectivity regions are presented for the whole cohort, term-born cohort, and preterm-born cohort (left to right) across composite, externalizing, and internalizing CBCL scores (top to bottom). Only ROIs with a degree ≥ one-sixth of the highest ROI are displayed; node size is proportional to degree. The 40% coverage in figure 3F indicates the last coverage level at which independent validation remained significantly above chance; coverage levels above 40% are not significant.

**Figure 4.**
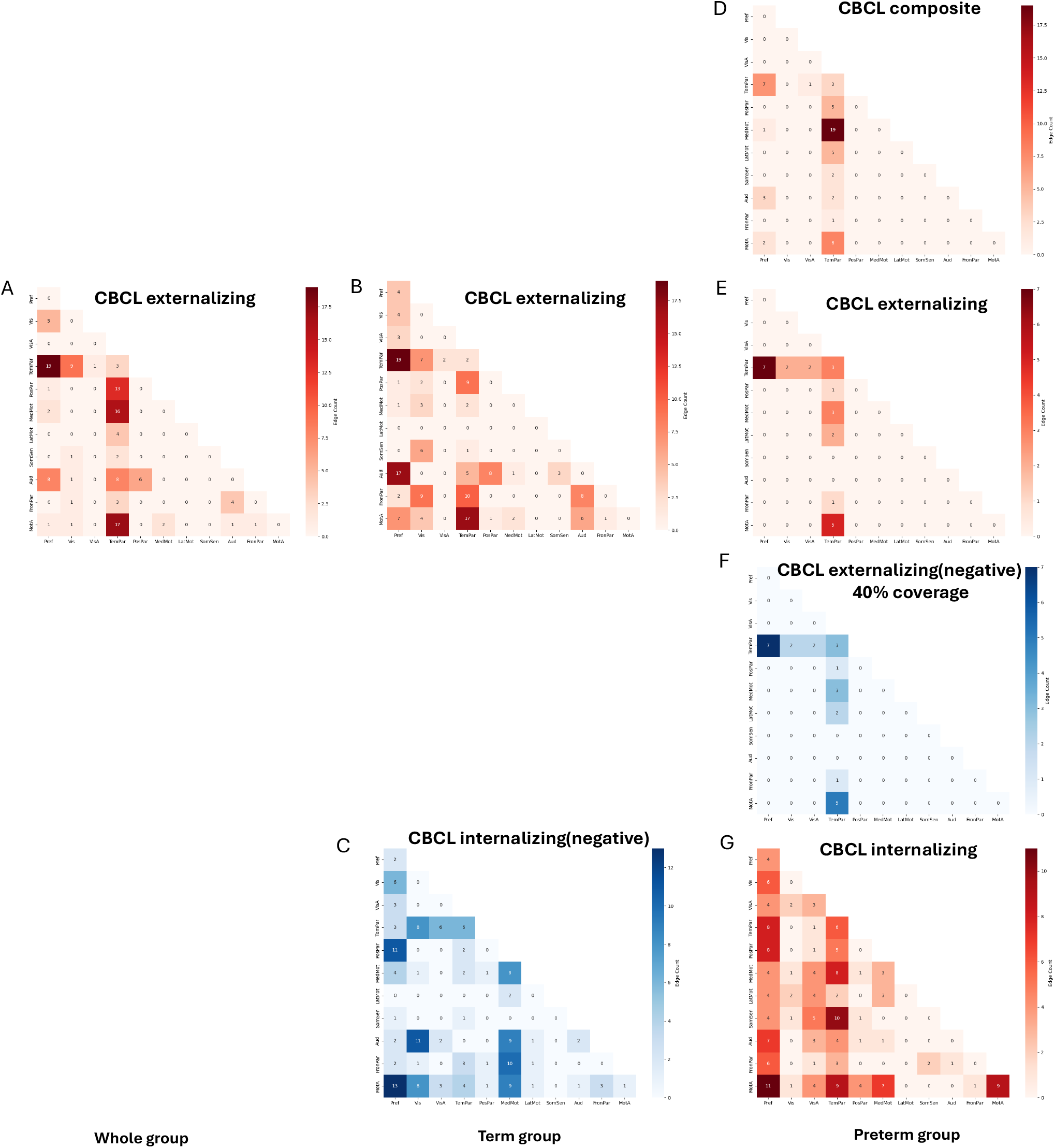
Predictive networks for CBCL across cohorts and subscales. Connections plotted as the number of edges within and between each pair of canonical networks for the whole cohort, term-born cohort, and preterm-born cohort respectively(left to right) across externalizing, and internalizing CBCL scores (top to bottom). The 40% coverage in figure 4F indicates the last coverage level at which independent validation remained significantly above chance; coverage levels above 40% are not significant.

For the term-born group, the CBCL prediction models were statistically significant for the CBCL externalizing score (see Table 2). Nodes were primarily located in right insula lobe and left lingual gyrus (Figure 3B). The prefrontal network, temporoparietal network and auditory network are prominent in CBCL externalizing model, especially the connection between prefrontal network and temporoparietal network (Figure 4B). In addition, for the term group, a negative network associated with the CBCL internalizing score was identified (see Table 2). Nodes within this network were primarily located in the left insula lobe and right mid orbital gyrus (Figure 3C). The prefrontal, medial motor and motor association networks were core part of the prediction model, with prominent connections observed between prefrontal network and motor association network (Figure 4C).

For the preterm-born group, the CBCL prediction model was statistically significant for CBCL composite score, CBCL externalizing score and CBCL internalizing score (see table 2). Nodes in CBCL composite model and CBCL externalizing model were mainly located in left fusiform gyrus and left parahippocampal gyrus (Figure 3D-3E), while nodes of CBCL internalizing model are primarily in right rectal gyrus, right parahippocampal gyrus and left middle temporal gyrus (Figure 3G). Temporoparietal network is dominative in CBCL composite model and CBCL externalizing model (Figure 4D-4E), while prefrontal, temporoparietal and motor association networks are actively involved in CBCL internalizing model (Figure 4G). In addition, for the preterm group, a negative network associated with the CBCL externalizing score was identified (40% coverage) (see Table 2). Node with highest degree was right middle temporal gyrus (Figure 3F). The temporoparietal network is the most prominent, especially the connection between temporoparietal network and prefrontal network (Figure 4F).

The ECBQ prediction model with the whole cohort was statistically significant for surgency score and negative affect score (see table 2). Nodes in right parahippocampal gyrus and left middle temporal gyrus were prominent in the surgency model (Figure 5A), whereas nodes in left and right inferior frontal gyrus were predominant in the negative affect model (Figure 5B). Accordingly, temporoparietal and motor association networks are highly engaged in the surgency model, especially the connection between the two (Figure 6A), while prefrontal, temporoparietal, motor association and auditory networks are actively involved, especially the connection between prefrontal network and temporoparietal network (Figure 6B).

**Figure 5.**
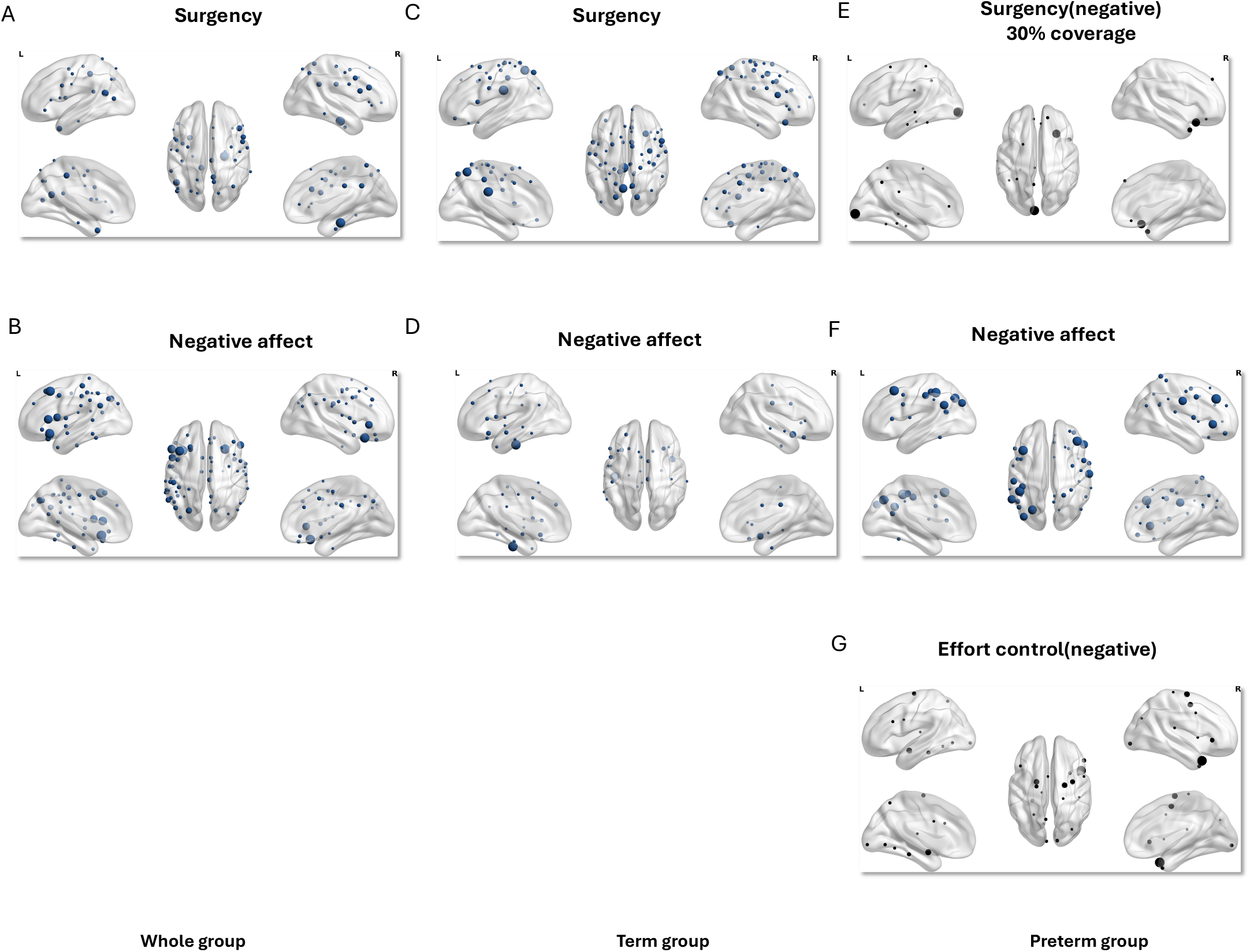
Predictive ROIs for ECBQ: high-degree regions across cohorts and subscales. High-connectivity regions are presented for the whole cohort, term-born cohort, and preterm-born cohort (left to right) across surgency, negative affect, and effort control scores (top to bottom). Only ROIs with a degree ≥ one-sixth of the highest ROI are displayed; node size is proportional to degree. The 30% coverage in figure 5E indicates the last coverage level at which independent validation remained significantly above chance; coverage levels above 30% are not significant.

**Figure 6.**
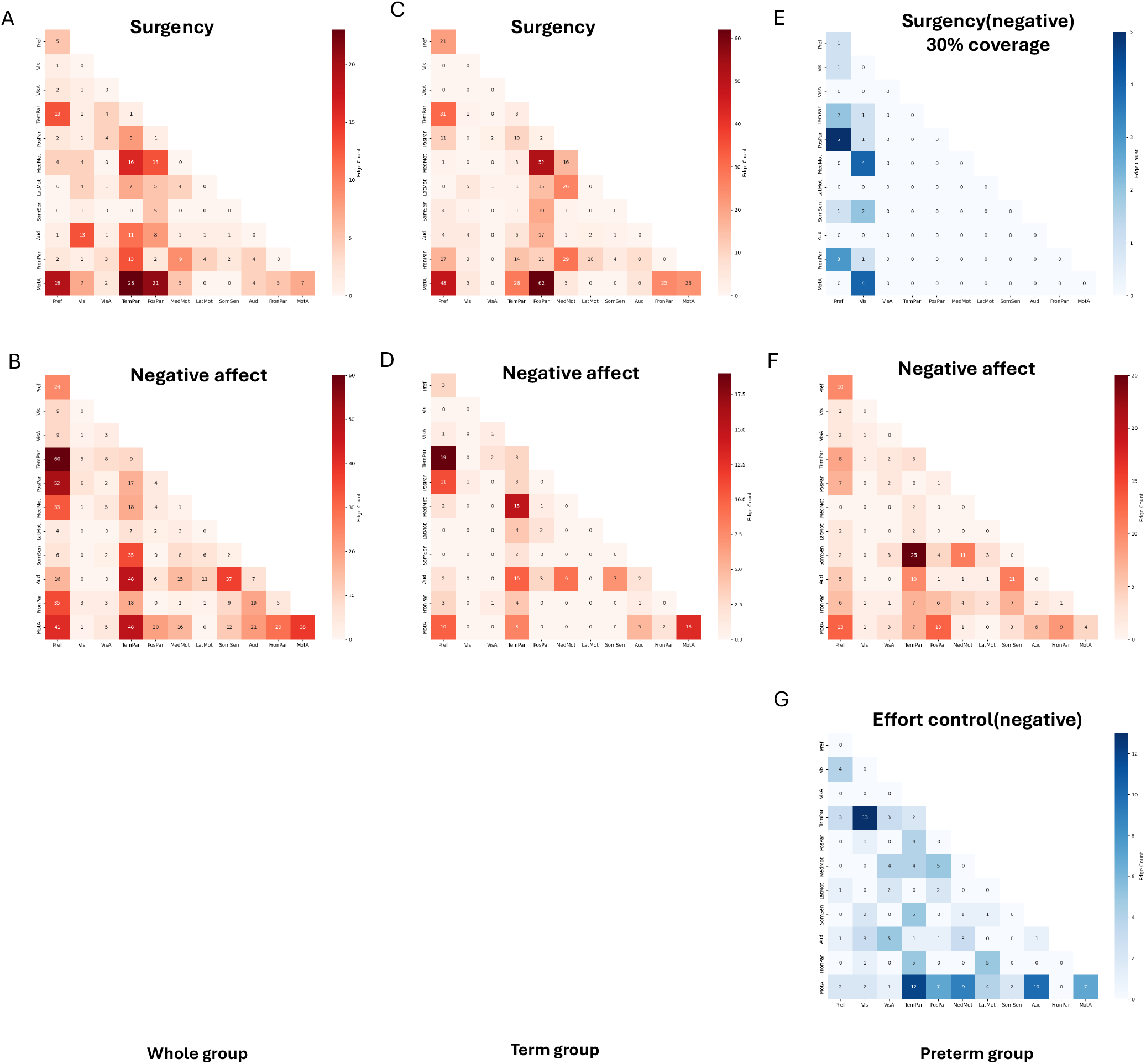
Predictive networks for ECBQ across cohorts and subscales. Connections plotted as the number of edges within and between each pair of canonical networks for the whole cohort, term-born cohort, and preterm-born cohort (left to right) across composite, fine, and gross motor scores (top to bottom). The 30% coverage in figure 6E indicates the last coverage level at which independent validation remained significantly above chance; coverage levels above 30% are not significant.

In the term-born group, the ECBQ model was statistically significant for surgency score and negative affect score (see table 2). Nodes were mainly in left posterior cingulate cortex and left precuneus in the surgency model (Figure 5C) while in the negative affect model, the node with highest connection is left inferior temporal gyrus (Figure 5D). The connectivity between postparietal network and motor assocition network played a dominant role in the surgency model, especially the connection between the two (Figure 6C), while the connectivity between prefrontal network and temporoparietal network was most prominent in the negative affect network, especially the connection between the two (Figure 6D).

In the preterm-born group, the ECBQ prediction model was statistically significant for the negative affect score (see Table 2). Nodes are mainly in left inferior parietal lobule, left middle frontal gyrus and right middle frontal gyrus (Figure 5F). The temporoparietal and motor association networks were highly active in negative affect model, with strongest connection observed between temporoparietal network and somatosensory network (Figure 6F). In addition, for the preterm group, two negative networks associated with surgency score (30% coverage) and effort control score respectively were identified (see Table 2). Nodes within surgency model were primarily located in left calcarine gyrus and right inferior frontal gyrus (Figure 5E), while nodes within effort control model were primarily located in right medial temporal pole and right superior frontal gyrus (Figure 5G). The prefrontal and visual networks were core part of the surgency model (Figure 6E), while temporoparietal and motor association networks were dominant in the effort control model (Figure 6G).

## Discussion

This study shows that neonatal resting-state functional connectivity predicts multiple socioemotional and behavioral outcomes at 18 months using a stability-driven ROI-constrained connectome-based predictive modeling framework. Significant models were identified for CBCL and ECBQ dimensions across the whole cohort, term-born infants, and preterm-born infants. These findings suggest that individual differences in externalizing behavior, internalizing behavior, surgency, negative affect, and effortful control are already reflected in large-scale neonatal functional organization. Importantly, these results should not be interpreted as deterministic prediction, but rather as evidence that early functional connectivity contains meaningful information related to later behavioral and temperamental variation.

Across analyses, prefrontal and temporoparietal systems were consistently implicated, particularly in models predicting CBCL externalizing and ECBQ negative affect. In the whole cohort, CBCL externalizing was characterized by strong temporoparietal involvement and prominent connections between temporoparietal and prefrontal networks. Similar patterns were observed in the term-born externalizing model and in the preterm-born negative externalizing network. For ECBQ negative affect, prefrontal–temporoparietal connectivity was also prominent in both the whole cohort and term-born group. These findings are consistent with prior work showing that infant large-scale functional connectivity is associated with affective and behavioral differences, including negative affectivity and later behavioral outcomes (Kelsey et al., 2021; Ravi et al., 2023).

The present results also extend previous neonatal studies that have focused mainly on predefined emotion-related circuits, especially amygdala-centered pathways. Neonatal amygdala connectivity has been associated with fear, negative reactive temperament, internalizing symptoms, and early-emerging callous-unemotional traits (Thomas et al., 2019; Filippi et al., 2021; Rogers et al., 2017; Brady et al., 2024). Our findings are compatible with this literature, but suggest that predictive information is distributed across broader systems, including prefrontal, temporoparietal, medial temporal, sensory-association, and motor-association networks. This broader organization is plausible because early socioemotional traits likely depend on the coordination of systems involved in attention, salience processing, sensory integration, emotional reactivity, and regulation.

Several models also involved medial temporal, fusiform, parahippocampal, and orbitofrontal-related regions. In the whole cohort, CBCL externalizing was dominated by the left fusiform gyrus and right medial temporal pole. In preterm-born infants, CBCL composite and externalizing models involved the left fusiform and left parahippocampal gyri, whereas CBCL internalizing involved rectal, parahippocampal, and middle temporal regions. ECBQ surgency in the whole cohort also involved parahippocampal and temporal nodes. These findings suggest that later socioemotional outcomes are not only related to higher-order networks, but also to regions involved in social-perceptual processing, contextual integration, salience, and affective valuation. This interpretation is broadly consistent with evidence that neonatal cerebellar structure and functional connectivity predict later social and emotional development (Kim et al., 2024).

A key contribution of this study is the comparison between term-born and preterm-born infants. Although both groups showed significant predictive models, the implicated connectivity patterns differed. In term-born infants, predictive models more often involved prefrontal–temporoparietal connectivity, auditory participation, insular or lingual regions, and posterior midline regions such as posterior cingulate cortex and precuneus. In preterm-born infants, models more often emphasized temporoparietal, motor-association, medial temporal, parahippocampal, and frontal systems, as well as negative networks for externalizing, surgency, and effortful control. These differences suggest that preterm birth may alter which neonatal functional systems are most relevant to later behavioral and temperamental outcomes.

This cohort divergence is developmentally plausible. Late gestation is a period of rapid development in long-range connectivity and inter-network coordination. For preterm-born infants, part of this period occurs in the extrauterine environment, which may alter sensory input, autonomic regulation, stress physiology, and other developmental experiences during a sensitive window. Therefore, the connectivity patterns associated with later behavior in preterm-born infants may reflect a partly different developmental pathway rather than simply a weaker version of term-born development. This interpretation is consistent with evidence that very preterm children show atypical resting-state connectivity and that third-trimester extrauterine exposure alters central autonomic network development (Kozhemiako et al., 2019; Christoffel et al., 2025).

More broadly, neonatal functional connectivity may reflect multiple prenatal and perinatal influences. The intrauterine environment is shaped by maternal biological and environmental factors, including stress physiology, cortisol exposure, depressive symptoms, and social adversity. Prior work has linked neonatal connectivity to maternal cortisol, prenatal stress, maternal depression, social disadvantage, and later behavioral outcomes (Graham et al., 2019; Marr et al., 2023; Soe et al., 2016; Ramphal et al., 2020; Herzberg et al., 2024). These influences may partly contribute to the predictive signal captured by neonatal connectome-based models. However, because the present study did not directly model detailed parental, family, or environmental factors, the observed brain–behavior associations should not be interpreted as purely neural or causal.

Methodologically, this study extends CPM by using a ROI-constrained strategy. Here, stability-driven refers to identifying edges and ROIs that are repeatedly selected across many cross-validation folds and repetitions, rather than relying on features selected from a single split. This approach prioritizes reproducible high-degree ROIs and reduces the influence of unstable edges. The signal-to-noise analysis supported this logic: prediction accuracy declined as progressively lower-degree ROIs were added, and this decline closely tracked a reduction in signal-to-noise ratio. At the same time, independent validation showed that lower-degree signal edges can still contribute predictive information. Thus, ROI-constrained CPM provides a compromise between full connectome-wide prediction and narrowly predefined circuit analyses.

Several limitations should be noted. First, although several models were statistically significant, effect sizes were modest and varied across outcomes and cohorts, as expected for complex early behavioral phenotypes. Second, CBCL and ECBQ scores were based on caregiver report, which is developmentally appropriate but may not capture all aspects of infant behavior. Third, although whole-brain modeling can identify distributed predictive patterns, it provides less mechanistic specificity than targeted circuit analyses. Future studies should combine whole-brain predictive modeling with focused tests of limbic, cerebellar, autonomic, and sensory systems, and should include richer measures of prenatal, family, and environmental context.

In conclusion, neonatal resting-state functional connectivity contains information associated with socioemotional and behavioral outcomes at 18 months. The findings highlight distributed large-scale systems, especially prefrontal, temporoparietal, medial temporal, and sensory-association networks, and show that term-born and preterm-born infants differ in the connectivity patterns associated with later behavior. These results support a developmental view in which early functional brain organization reflects both biological maturation and environmental influences, contributing to later individual differences in socioemotional development.

## Supporting information

see Supplementary Table 1

## Acknowledgement

All calculations were performed on the high performance cluster maintained by the Trinity College Research IT unit. The research was supported by funding through a joint scholarship programme of the China Scholarship Council and Trinity College Dublin. Data used in the preparation of this manuscript were obtained from the National Institute of Mental Health (NIMH) Data Archive (NDA). NDA is a collaborative informatics system created by the National Institutes of Health to provide a national resource to support and accelerate research in mental health. Dataset identifier(s): <u>10.15154/f6j3-vx21</u>. This manuscript reflects the views of the authors and may not reflect the opinions or views of the NIH or of the Submitters submitting original data to NDA.

## Author Contributions

Arun L. W. Bokde conceptualized the study, supervised the project and revised manuscript. Mi Zou developed the innovative methodology, performed the data analysis, and wrote and revised the manuscript.

