## Supplementary material for "Neonatal Resting-State Functional Connectivity Predicts Socioemotional and Behavioral Outcomes at 18 Months": see Supplementary Table 1

### Table 1 Correlation between prediction accuracy and SNR

| | Whole cohort | Term cohort | Preterm cohort |
|---|---|---|---|
| CBCL composite | 0.94 | 0.86 | 0.71 |
| CBCL Internalizing | 0.81 | 0.83 | 0.63 |
| CBCL Externalizing | 0.89 | 0.94 | 0.66 |
| Surgency | 0.84 | 0.84 | 0.59 |
| Negative affect | 0.75 | 0.82 | 0.56 |
| Effort control | 0.57 | 0.90 | 0.84 |

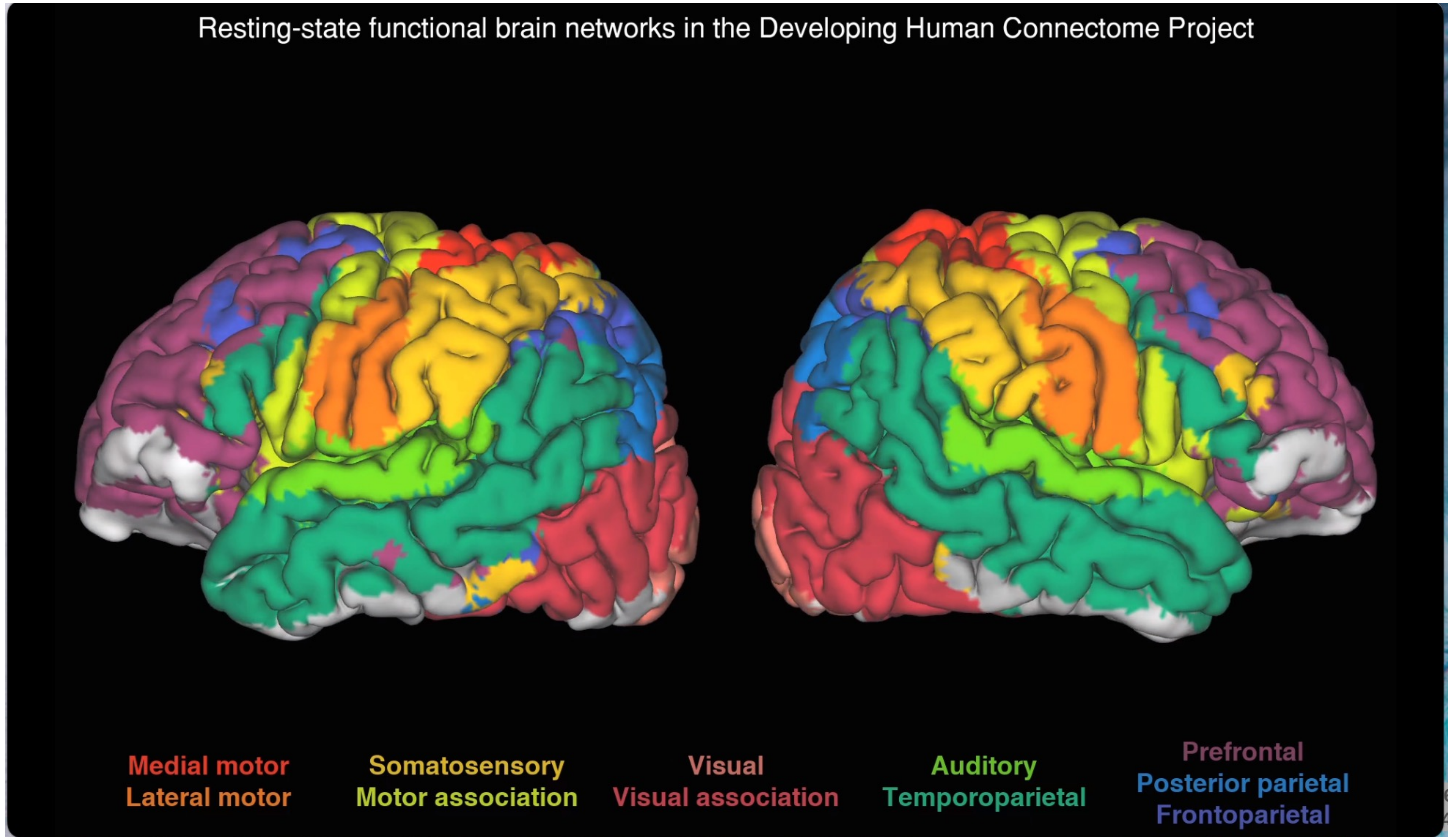


**Figure S1. Network parcellation and RSN assignment.**
Group-level independent component analysis defined canonical resting-state networks (term infants at 43.5–44.5 weeks postmenstrual age).
